# Mechanical characterization of soft biomaterials: which time and spatial scale to choose?

**DOI:** 10.1101/2024.04.30.591439

**Authors:** E.S. Krivega, S.L. Kotova, P.S. Timashev, Y.M. Efremov

## Abstract

The mechanical properties of soft gels hold significant relevance in biomedicine and biomaterial design, including the development of tissue engineering constructs and bioequivalents. It is important to adequately characterize the gel’s mechanical properties since they play a role both in the overall structural properties of the construct and in cells’ physiological responses. The question remains which approach for mechanical characterization is most suitable for specific biomaterials. Our investigation is centered on the comparison of three types of gels and four distinct mechanical testing techniques: shear rheology, compression, microindentation, and nanoindentation by atomic force microscopy. While analyzing an elastic homogeneous synthetic hydrogel (polyacrylamide gel), we observed close mechanical results across the different testing techniques. However, our findings revealed more distinct outcomes when assessing a highly viscoelastic gel (Ecoflex) and a heterogeneous biopolymer hydrogel (enzymatically crosslinked gelatin). To ensure the precise data interpretation, we introduced correction factors to account for the boundary conditions inherent in many of the testing methods. The results of the study underscore the critical significance of considering both the temporal and spatial scales in mechanical measurements of biomaterials. Furthermore, they encourage the employment of a combination of diverse testing techniques, particularly in the characterization of heterogeneous viscoelastic materials such as biological samples. The obtained results will contribute to refining mechanical testing protocols and advance the development of soft gels for tissue engineering.

## 1. Introduction

Designing materials for tissue engineered constructs and tissue substitutes is one of the main tasks of regenerative medicine (Chan and Leong, 2008; Chun et al., 2018; Hanczar et al., 2021; Stratakis, 2018). Mechanical properties of such materials are currently recognized as an important factor for the construction of successful replacement materials for damaged tissues and tissue engineered constructs (Chan and Leong, 2008; Hanczar et al., 2021; Matellan and Del Río Hernández, 2019; Roeder, 2013). Specifically, a replacement material should have mechanical properties sufficiently close to such of the host tissue at different spatial levels, the feature that is called “mechano-mimetic” (Cappitti et al., 2023; Kutty and Webb, 2009). At the macroscale, an implanted tissue engineered construct will need to provide the mechanical and shape stability to the tissue defect under physiological deformations and loads. Moreover, the mechanobiology studies showed importance of the micromechanical properties of a material for cells interacting with the scaffold. Mechanosensitivity, the sensation of the substrate stiffness and response via a change in the morphology and functional characteristics was demonstrated for many cell types, including stiffness-affected differentiation of mesenchymal stem cells (Engler et al., 2006; Lee et al., 2019). A potential construct material for soft tissue such as parenchymal organs, skin, and cartilage, therefore should have appropriate mechanics. Soft hydrogels based on natural and synthetic polymers are often used in cell mechanics and mechanobiology to represent soft tissues (Cappitti et al., 2023; Kutty and Webb, 2009; Palchesko et al., 2012; Vedadghavami et al., 2017; Markert et al., 2013).

While the mechanics of soft materials for the use in biomedical applications have been extensively studied, several aspects still remain that require attention. For example, several mechanical testing techniques that are available for soft materials often provide inconsistent data (Di Lorenzo et al., 2015; McCarthy et al., 2023; Navindaran et al., 2023; Buffinton et al., 2015; McKee et al., 2011). The discrepancies can be caused by different lengths, deformations, and time scales applied by different techniques, and by a different behavior of the tested material on these scales. The latter includes surface effects, such as adhesion and surface energy, nonlinearity, heterogeneity, viscoelasticity, and poroelasticity among others (Di Lorenzo et al., 2015; Megone et al., 2018; Navindaran et al., 2023). The testing techniques themselves may contain specific sources of error, often related to the sample preparation and boundary conditions. Therefore, the comparison of different mechanical testing methods for soft materials is required to establish the specificity of each technique for proper material assessment and to further translate these measurement techniques to soft biological tissues, where validation of such techniques is also required.

Here, we applied four different mechanical testing techniques for the characterization of three types of soft gels. These techniques included shear rheology, compression, microindentation, and nanoindentation by atomic force microscopy (AFM), covering different spatial and time experimental scales. The selected materials included a polyacrylamide gel (PAAG), which is a hydrogel formed by a highly homogenous synthetic polymer mesh and is extensively used in mechanobiology studies (Denisin and Pruitt, 2016; Grevesse et al., 2014), an enzymatically crosslinked gelatin hydrogel (Broderick et al., 2005; Liu et al., 2020), representing the class of biological polymers, and Ecoflex^TM^, representing the class of soft biocompatible silicone gels, which, unlike the previous two materials is not a hydrogel, but has a comparable stiffness and is used in bioelectronics and wearable sensors (Kim et al., 2023; Liao et al., 2020; Mansy et al., 2008). We compared the measured mechanical parameters considering the peculiarities of each method and each sample type. Special attention was given to frequency-dependent properties since the viscoelastic behavior is common to many biological tissues. For that, the experiments were conducted at different measurement rates, and effective frequencies were calculated for the used methods where frequency is not prescribed by default.

## 2. Material and Methods

### 2.1 Gels preparation

Polyacrylamide gels (PAAGs) were prepared from solutions of acrylamide and N,N′-methylene bisacrylamide (Sigma) in Milli-Q water with 10% ammonium persulfate (Sigma) and N,N,N’,N’-Tetramethylethylenediamine (Fluka, Germany). Final concentrations of acrylamide and bis-acrylamide were 5% and 0.3% w/v, respectively. For AFM experiments, a total solution volume of 100 µL was polymerized between clean glass square coverslips (Menzel–Gläser, Germany) with 100-µm thick spacers at the edges (Parafilm M, Sigma Aldrich). After 10 min of incubation, the top coverslip was removed and the sample was extensively washed with Milli-Q water. For the rheometric measurements, PAAG was polymerized directly between the plates of the rheometer in a plate-plate configuration (25 mm diameter of the upper plate) with a 1 mm gap between the plates (0.5 mL volume of the solution).

Ecoflex 00-10 (Smooth-On, Easton, PA, USA), where 00-10 is the Shore hardness, was used as an example of a soft silicone gel, formed from a two-component platinum-catalyzed room-temperature curable silicone. As per manufacturer instructions, two liquid components of the Ecoflex were mixed and stirred manually with a 1:1 ratio by mass. The mixture was then poured into a customized mold or onto the surface of a Petri dish, degassed in a vacuum chamber to remove trapped air bubbles, and left overnight to form a gel with a thickness of 2-4 mm for rheology, compression, and microindentation experiments or 100-200 µm for AFM experiments.

Gelatin from porcine skin, Type A, ∼300 g Bloom (Sigma) was used at a concentration of 10% in water and crosslinked by the addition of microbial transglutaminase (mTG) (100 U/g, TD Biopreparat, Moscow, Russia). A 10% solution of mTG in PBS was prepared and filtered through a 0.45 μm syringe filter (Agilent Technologies). The final concentration of mTG was 10 U/g (enzyme/gelatin). The cross-linking reaction was conducted at 37°C in a humid environment for 10 hours, after which the gel was extensively washed with PBS. The gels were prepared in the shape of discs with the diameter of 25 mm and thickness of 2-4 mm or films with the thickness of 100-200 µm for AFM experiments.

### 2.2 Microindentation

Microindentation testing was performed with a Mach-1™ v500csst universal micromechanical system (Biomomentum Inc., Laval, QC, Canada). A metal spherical indenter with a radius of 3.175 mm was used. Before the measurements, the exact position of the substrate (metal plate) was found using the instrument’s “Find Contact” function. This function ensures that the indenter moves until the specified load is registered on the force sensor. Subsequently, the sample thickness was calculated relative to the vertical position of the substrate. Using the same function (“Find Contact”), the sample was indented to a certain load, adjusted in a way to ensure an indentation depth of the order of a third of the sample thickness or lower. The indentations were applied in 9 separated points on a grid with a size of 3×3 mm and an indentation map was obtained for each of the applied indentation speeds, which were 0.1, 0.2, 0.4, 0.8, and 1.6 mm/s. The results of this testing process are force-distance (F-Z) curves, numerical processing of which was performed using Python scripts (https://github.com/yu-efremov/ViscoIndent, accessed on 1 February 2024) developed in the previous works (Efremov et al., 2019, 2017) using the Hertz’s and Ting’s models (Ting, 1966) for the elastic and viscoelastic processing, respectively. Elastic processing was performed with the Hertz’s model with the correction of the sample’s finite thickness:

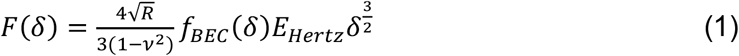

where *F* is the force acting on the cantilever tip; *δ* is the indentation depth; *ν* is the Poisson’s ratio of the sample (assumed to be time-independent and equal to 0.5); *R* is the radius of the indenter; *E*_*Hertz*_ is the Young’s modulus with the assumptions of Hertz’s theory (“apparent” indentation modulus), and *f*_*BEC*_*(δ)* is the bottom-effect correction. The latter is a multiplicative analytically derived correction for the indentation contact models that accounts for the finite sample thickness (Efremov et al., 2019, 2017). The multiplicative coefficients were taken from the work (Garcia and Garcia, 2018). When a significant hysteresis indicating viscoelastic effects was presented in the force curves, they were processed using the viscoelastic model:

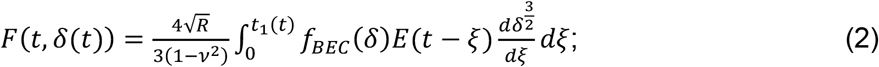

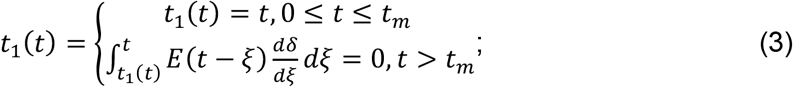

where *t* is the time initiated at contact; *t*_*m*_ is the duration of the approach phase; *t*_*1*_ is the auxiliary function determined by Equation (3); *ξ* is the dummy time variable required for the integration; and *E(t)* is Young’s relaxation modulus for the selected rheology model. Here, we used the power-law rheology (PLR) model (a fractional element, springpot, in parallel with a spring) (Kollmannsberger and Fabry, 2009):

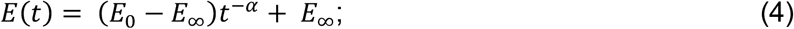

where *E*_0_ is the relaxation modulus at *t* = 1 s (scale factor of the relaxation modulus); *α* is the power-law exponent; *E*_∞_ is the long-term modulus. The selection of a particular viscoelastic model does not play a significant role, since the extracted parameters of the storage and loss Young’s moduli were calculated, which do not depend on the selected viscoelastic model as shown in the previous study (Efremov et al., 2019).

### 2.3 Atomic force microscopy (nanoindentation)

The mechanical properties at the nanoscale were measured with a Bioscope Resolve atomic force microscope (Bruker, Santa Barbara, Goleta, CA, USA) mounted on an Axio Observer inverted fluorescence microscope (Carl Zeiss, Germany). PeakForce QNM-Live Cell probes (PFQNM-LC-A-CAL, Bruker AFM Probes, Camarillo, CA, USA) with a pre-calibrated spring constant (average value of 0.1 N/m) and tip radius (70 nm) were used, and the deflection sensitivity (nm/V) was calibrated from the thermal spectrum using the value of the spring constant. Experiments were conducted in a liquid environment that was Milli-Q water with the addition of a detergent (0.01% Triton X-100, Sigma) to reduce the adhesion between the probe and the sample surface. Nanomechanical and topography maps were acquired in the force–volume mode with the map sizes from 80 × 80 to 10 × 10 µm and from 32 × 32 to 128 × 128 point measurements. The force curves (*F-Z* curves) had a vertical ramp distance of 3 μm, a vertical piezo speed in a range of 1 to 200 μm/s, and a trigger force of 1.5–2.5 nN (depending on the sample studied). The typical indentation depth was 200–400 nm. Examples of the force curves together with the model fits are presented in Figure 1.

**Figure 1.**
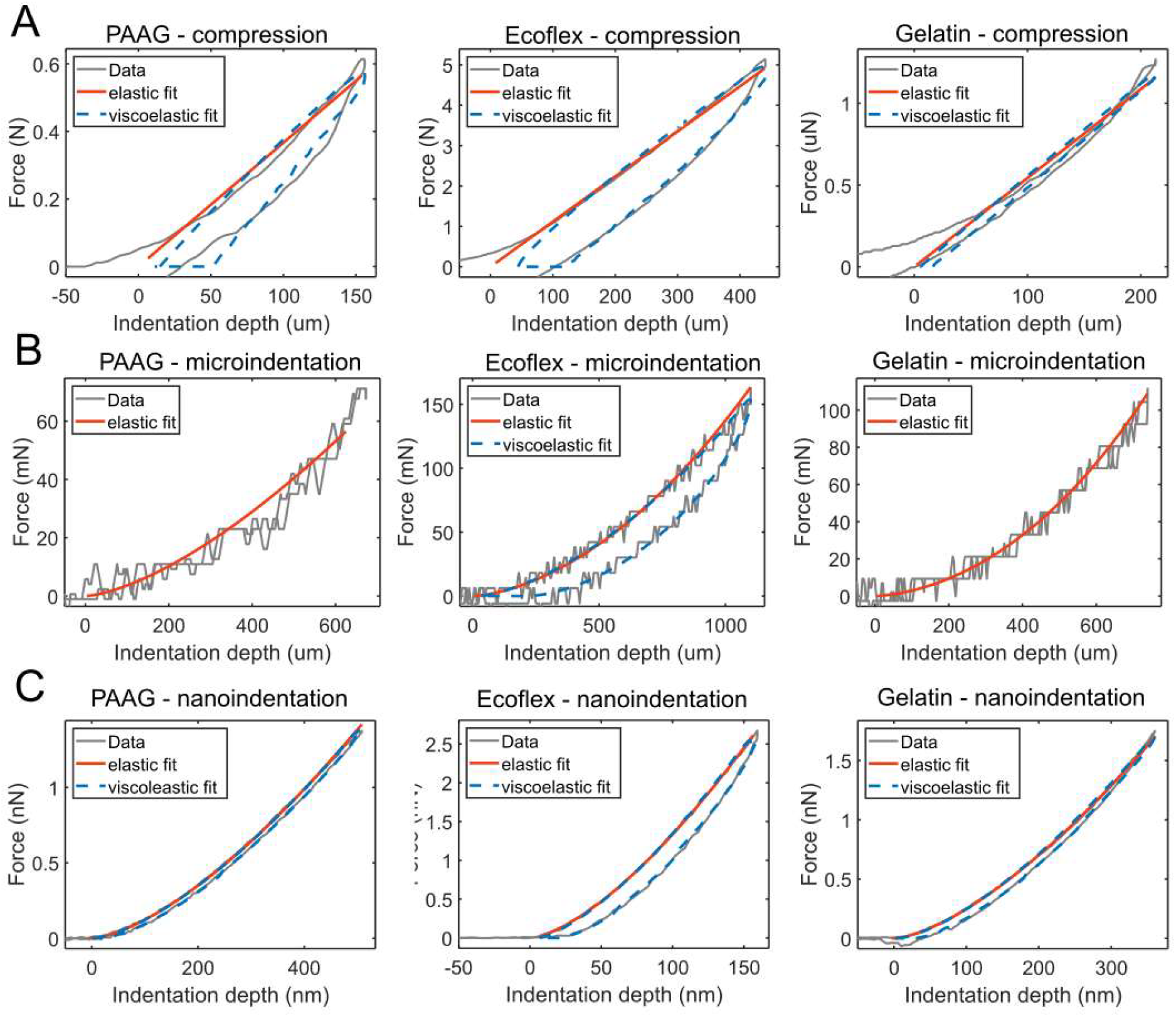
Examples of experimental data with the model fits (elastic and viscoelastic where applicable) obtained on polyacrylamide gels (PAAG), Ecoflex gels, and gelatin gels crosslinked with mTG for different techniques: (A) compression; (B) microindentation (C) nanoindentation by AFM. A more pronounced hysteresis in loading cycles corresponds to a larger degree of viscoelasticity, as can be seen for the Ecoflex sample in all the experiments and for the compression of a PAAG sample.

In the microindentation experiments, the Hertz’s and Ting’s models (Equations 1-4) were used for the processing with the same numerical procedures. The bottom-effect correction was not used since the indentation depth and tip radius (hundreds of nm) were much smaller than the sample thickness (100-200 µm).

### 2.4 Compression

The compression experiments were performed with a Mach-1™ v500csst universal micromechanical system (Biomomentum Inc., Laval, QC, Canada) on 25 mm-diameter disks cut from each type of hydrogels. Before the measurements, the zero distance between the metal plates was found using the instrument’s “Find Contact” function. Subsequently, the disk thickness (2-4 mm) was calculated relative to the vertical position of the bottom plate. During the measurements, the loading force was chosen so that the compression of the sample was approximately 30% of the thickness. After achieving the specified force, the upper plate moved up with the same speed to record a hysteresis in the loading cycle. The upper plate movement speeds were 0.1, 0.2, 0.4, 0.8, 1.6 mm/s. The elastic processing (effective Young’s modulus) was performed based on the slope of the force (F) versus the nominal compressive strain (ε) curve and geometric parameters of the sample:

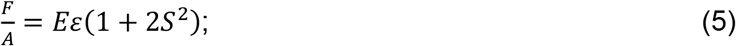

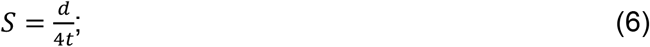

where *d* is the disk diameter, *w* is the disk width, *A* is the area of one loaded surface. *S* is the shape factor defined as the ratio of one loaded area to the force-free area, it is only required for the experiments on Ecoflex gel, where strong bondage of the gel surfaces to the metal plates was observed (Yeoh et al., 2002). The viscoelastic processing was performed for the selected samples using the Ting’s model modified for the case of a cylinder compression.

### 2.5 Rheology

The rheological properties of the samples were characterized with a Physica MCR 302 rheometer (Anton Paar, Austria) with a plate-plate geometry (25 mm diameter) at a controlled temperature of 25°C. Samples in the shape of disks with the diameter of 25 mm and thickness of 1-2 mm were tested. The oscillatory strain sweep test was performed at a fixed frequency of 1 Hz to determine the linear viscoelastic region (LVR) of each type of a gel, and the frequency sweep test was performed for the viscoelastic characterization. The tests were repeated at least three times, the storage modulus, G’ (Pa), and the loss modulus, G’’ (Pa), were measured. For the strain sweep test, 20 points were chosen with a range between 0.01 - 10% of the sweep amplitude. The frequency sweep test was then performed at a 0.1% strain amplitude that was within the LVR of each gel type; 25 points were chosen within the logarithmically spaced frequency range between 0.01 - 100 Hz. The Young’s modulus (E) was calculated from the shear modulus (G) with the following equation:

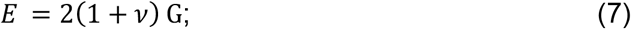

where *v* is the Poisson’s ratio, with the value of 0.48 for Ecoflex (Le et al., 2019), 0.48 for PAAG (Boudou et al., 2006), and 0.45 for gelatin gels (Van Otterloo and Cruden, 2016).

## 3. Results

### 3.1 Viscoelasticity in different mechanical experiments

Among the four mechanical measurement techniques applied here, for rheology, the shear moduli (shear storage and loss modulus vs frequency) were taken directly from the rheometer software, while for other methods additional processing steps to extract the mechanical parameters were performed. In the case of the indentation techniques (microindentation and AFM), the procedures for elastic and viscoelastic processing were previously developed. For the compression experiments, here we applied the viscoelastic processing similar to that for indentation with a cylindrical indenter by modifying the indentation model for the compression case. Examples of typical experimental force versus indentation (compression) curves are presented in Fig. 1. The presence of a hysteresis in the experimental (loading-unloading) cycle indicated the viscoelastic behavior and was most pronounced for Ecoflex samples. The set of corrections was applied for different methods and samples, as described below, including the finite thickness correction for microindentation experiments on all samples, and the correction for surface bonding for the Ecoflex gel compression.

The AFM technique also allowed characterization of the gels’ heterogeneity at the micrometer spatial scale (10×10 µm force volume maps). Examples of the force volume maps for each hydrogel that were acquired at the 180 µm/s piezo speed are presented in Fig. 2. The heterogeneity was assessed as the coefficient of variation (CV, the ratio of the standard deviation to the effective modulus average value, %) for the effective Young’s modulus acquired for such maps, and was similar for PAAG and Ecoflex gels (5-10%) and substantially larger for gelatin gels (15-25%).

**Figure 2.**
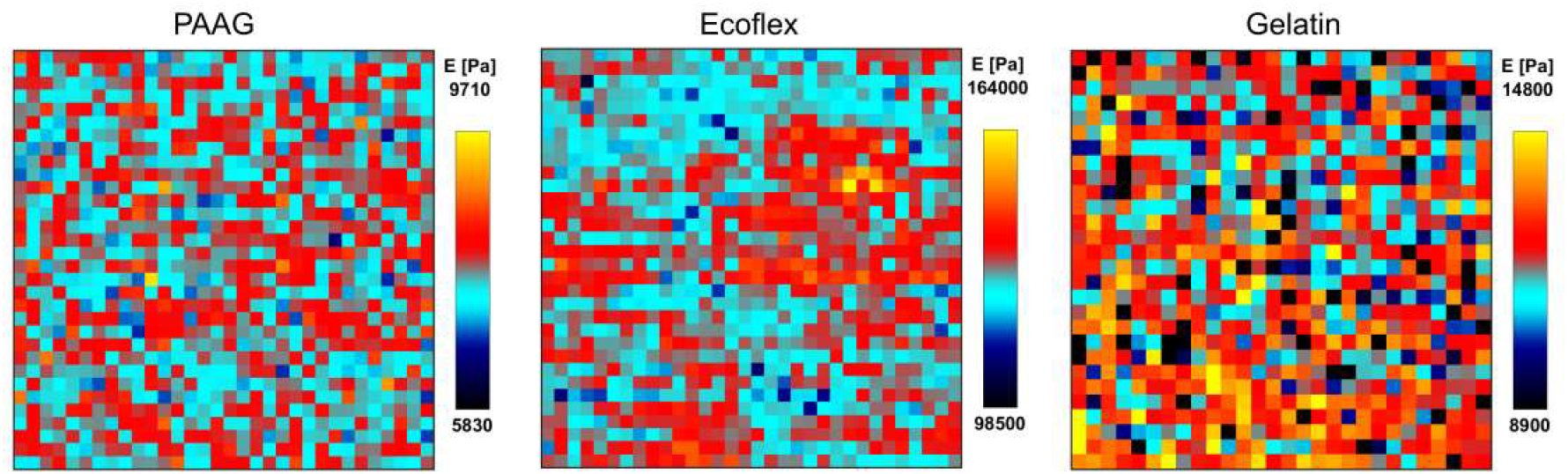
Examples of force volume maps (effective Young’s modulus) acquired by AFM for polyacrylamide gels (PAAG), Ecoflex gels, and gelatin gels crosslinked with mTG. Each map is an array of 32×32 indentations per 10×10 µm square area. The color bar for each map was set in the range of the mean modulus value ± 25% of the mean modulus value.

### 3.2 Characterization of PAAG samples

Mechanical properties of PAAGs were extensively characterized in the past years due to their widespread use in molecular biology, and, in particular, as substrates with adjustable mechanical properties for mechanobiology studies (Grevesse et al., 2014; Denisin and Pruitt, 2016)[10.3791/51010; 10.1021/acsami.5b09344]. Moreover, due to its well-defined elastic modulus, PAAG was used as a standardized sample for the comparison of AFM mechanical measurement results between different laboratories (Schillers et al., 2017). Although the behavior of PAAG is mostly elastic, poroelastic and viscoelastic effects can be pronounced at certain conditions of the gel preparation and measurements (Kalcioglu et al., 2012; Esteki et al., 2020).

In the current work, the prepared PAAG gels had mostly an elastic response in the rheological experiments, with the loss factor below 0.02 in most of the measured frequency range. At high frequencies, the glass transition was demonstrated in PAAG before (Abidine et al., 2015), and here an increase in the storage moduli was also noticed at frequencies above 10 Hz (Fig. 3).

**Figure 3.**
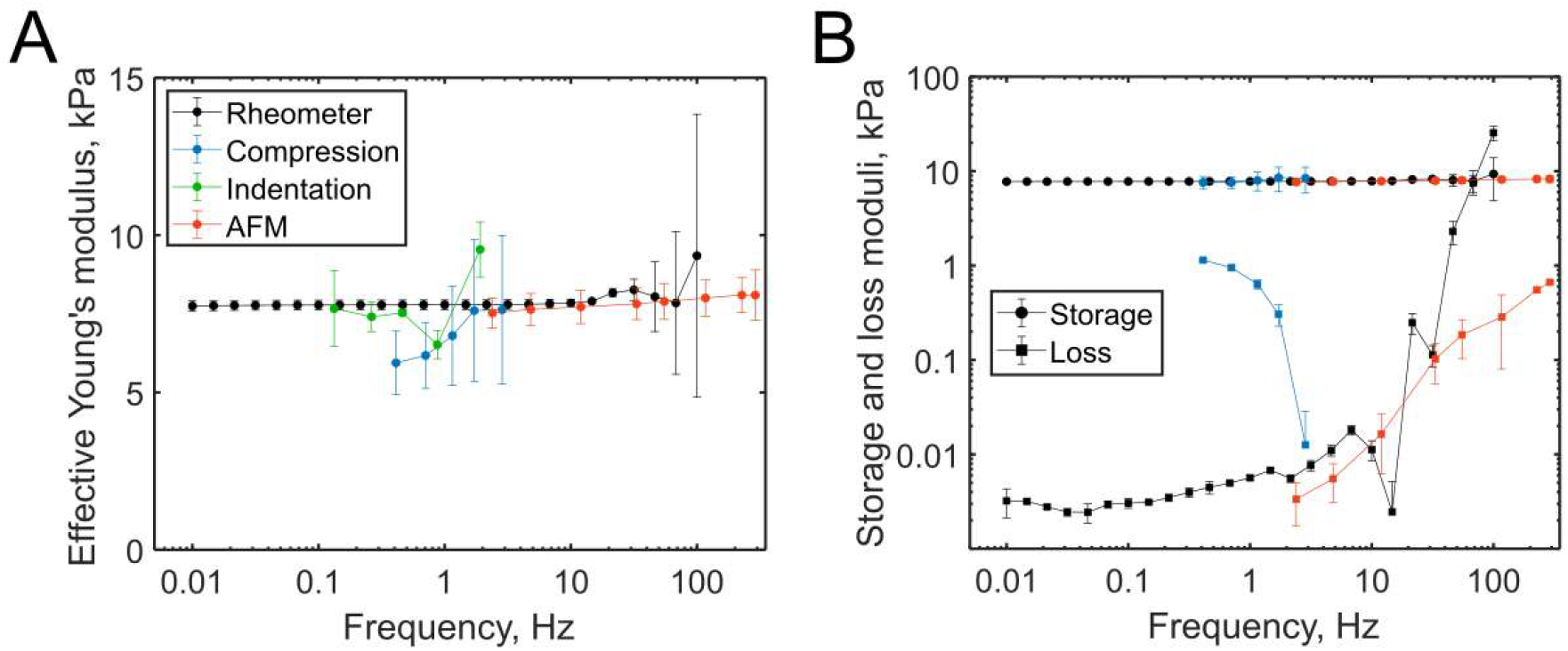
Comparison of five stiffness characterization methods for PAAG hydrogels. (A) Effective Young’s modulus versus effective frequency; (B) Storage and loss Young’s moduli versus effective frequency, microindentation data were not presented due to low hysteresis in the indentation cycle. The data are presented as mean±SD for measurements on three samples.

For other measurement techniques, the load speed was varied, and the effective frequency was calculated as an inverse of the contact time of each particular experiment. As shown in the previous studies, the introduction of “effective frequency” corresponding to the characteristic duration of an experiment can be used for the assessment of the complex Young’s modulus (de Sousa et al., 2017; Efremov et al., 2020a). The compression and microindentation were applied at low frequencies of ∼0.1-1 Hz due to the inertia in the large measurement system. On the other hand, the AFM indentation experiments were characterized by higher frequencies of ∼1-300 Hz, while low frequencies were limited by the drifts in the system. Microindentation experiments for PAAG and all other gels required correction for the finite thickness, since the probe (3.5 mm diameter) and the sample thickness (2-4 mm) were comparable. Such a type of correction is often used in AFM experiments and was applied here for microindentation (Dimitriadis et al., 2002; Garcia and Garcia, 2018). Notably, without the correction the measured modulus was ∼4 times higher. When aligned on the same graph, close values were acquired between the different techniques, in agreement with the previous studies on PAAGs (Abidine et al., 2015; Megone et al., 2018) and other hydrogels from synthetic polymers (Richbourg et al., 2022). The graphs for the effective elastic moduli are compared with the storage modulus measured by the rheometer on the logarithmic frequency scale and linear modulus scale, while the graphs with the storage and loss moduli are shown on the double logarithmic scale. The storage and loss moduli were characterized for all the applied techniques except microindentation, where the hysteresis in the approach-retraction cycle was too low for quantitative processing (<1%). The AFM-derived storage and loss moduli were relatively close to the rheology data, while the compression data showed a larger loss modulus. The latter fact could be due to energy losses resulting from the friction between the sample and compression plates, which contribute to the observed hysteresis in the loading cycle (Fig. 1A).

### 3.3 Characterization of Ecoflex gel samples

The same set of methods was applied to the Ecoflex silicone gel. Unlike PAAG and gelatin gels, Ecoflex is not a hydrogel, but a cross-linked silicone elastomer whose polymer network is swollen with silicone fluid (Curtis and Steichen, 2020). The gel was sticky to the surfaces, which allowed application of higher frequencies (up to 100 Hz) in rheology experiments without slippage. The silicone gel demonstrated high stickiness (adhesion) to the plates in the compression experiments as well, thus, an approximate solution for the analysis of bonded rubber blocks subjected to small compressions was applied here in the form of the multiplicative correction (Yeoh et al., 2002). Without this correction, the acquired mechanical parameters were greatly overestimated (4-5 times larger), while the addition of a lubricant (silicone oil) between the sample and compression plates led to values close to the ones acquired with the correction (Fig. 4A). At the same time, there was no much difference in the relative hysteresis area of the compression cycle with and without the lubricant (Fig. 4B). The AFM experiments were performed in a liquid (water) to remove capillary forces, reduce adhesion, and simulate conditions which are close to the other gel samples testing.

**Figure 4.**
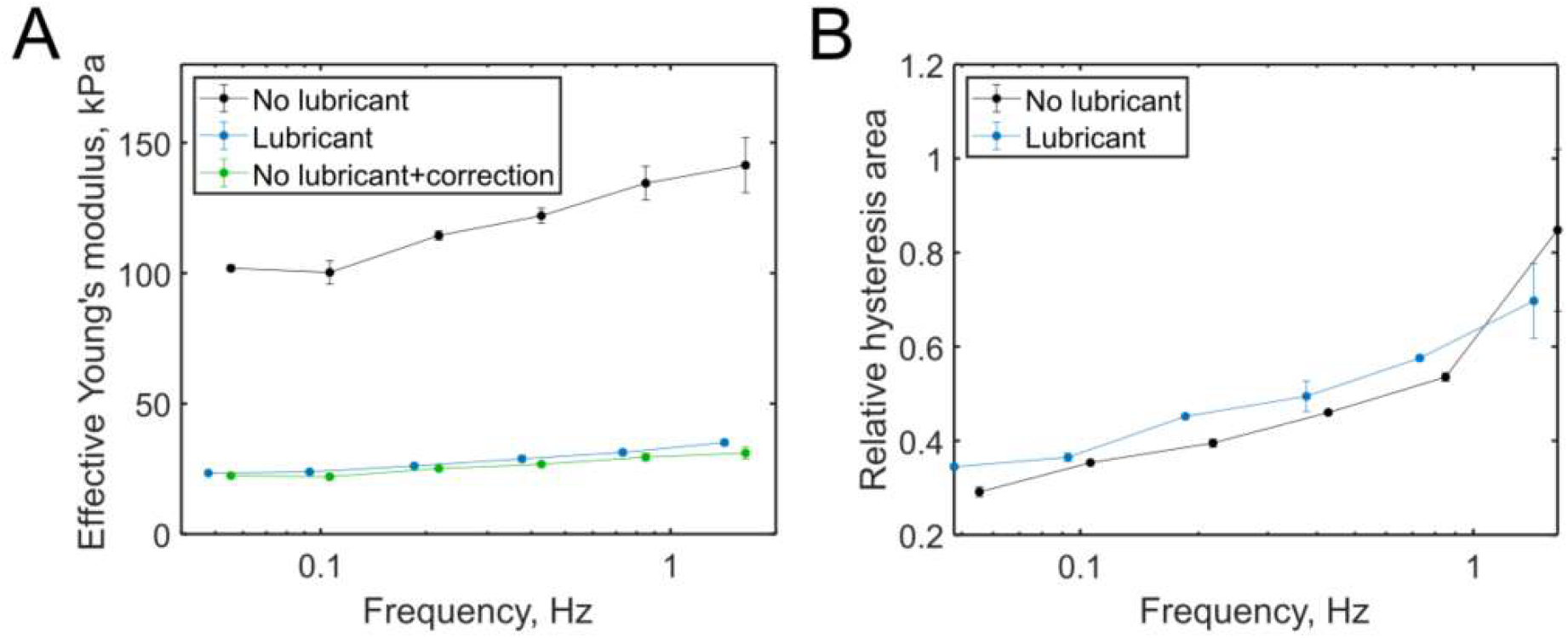
Compression of Ecoflex gels with and without the addition of a lubricant (silicon oil) between the sample and the plates. (A) Effective Young’s modulus versus effective frequency for the experiments with and without lubricant, and those without lubricant, but with applied correction for bonded surfaces; (B) Relative hysteresis area of the compression cycle for the experiments with and without lubricant. The data are presented as mean±SD for measurements on three samples.

Here we observed very pronounced viscoelastic behavior for the Ecoflex gel, in line with the previous studies, where a higher extent of viscoelasticity was observed for soft weakly crosslinked silicone gels (Kenry et al., 2015; Megone et al., 2018). There was a higher than fivefold increase in the storage modulus between the frequencies of 0.01 and 100 Hz in oscillatory rheology experiments (Fig. 5). This trend of an increase in the modulus with frequency was observed for all the applied measurement techniques within their own ranges of applied frequencies. The effective Young’s modulus values from the microindentation experiments and compression experiments were below the values acquired with the rheometer by 5-30%. The effective Young’s modulus from the AFM experiments was larger than that from the rheometer and demonstrated a weaker increase with frequency. The viscoelastic processing revealed similar dependencies for the storage and loss moduli between the rheology and microindentation data, while the loss moduli acquired by compression and AFM were substantially lower. In the compression experiments, the difference could stem from the system inertia and an imperfect shape of the silicone cylinders (e.g., non-flat sample surfaces). Higher storage modulus values and lower dissipation from the AFM data could originate from the higher degree of polymer crosslinking close to the surface as hypothesized before (Charitidis, 2011; Kenry et al., 2015).

**Figure 5.**
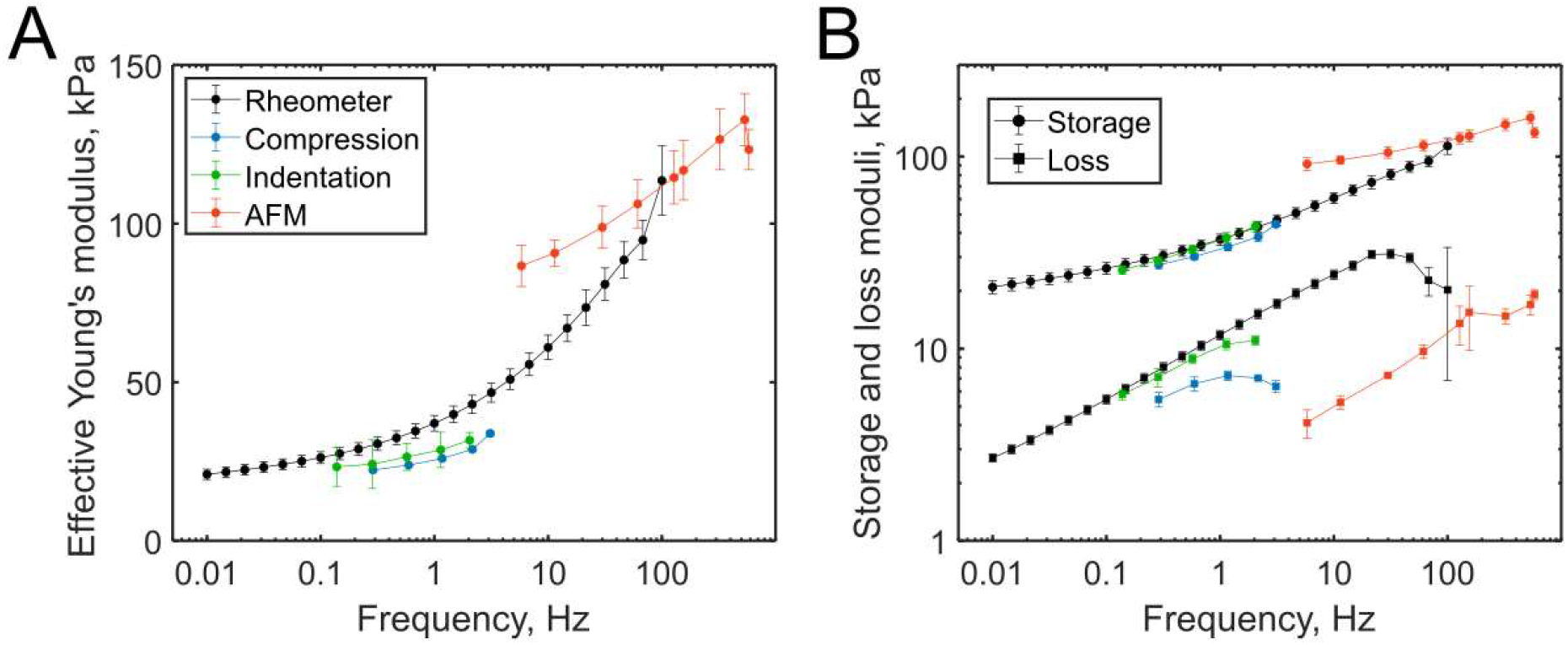
Comparison of five stiffness characterization methods for the Ecoflex gels. (A) Effective Young’s modulus versus effective frequency; (B) Storage and loss Young’s moduli versus effective frequency. The data are presented as mean±SD for measurements on three samples.

### 3.3 Characterization of gelatin samples

The gelatin hydrogels crosslinked with mTG represent an example of biological polymers (Broderick et al., 2005; Liu et al., 2020). The samples demonstrated a mostly elastic behavior in the rheological experiments, with the storage modulus almost independent of frequency (Fig. 6). Close mechanical data were obtained by microindentation and compression. The loss modulus measured in the compression experiments was larger compared to the rheology data, meaning higher energy losses, but was still rather low (several Pa). In the AFM experiments, the effective, storage, and loss moduli were higher than the corresponding data from the other methods. It has previously been shown for heterogeneous gels, that local moduli measured by AFM can be significantly larger than the global elastic modulus measured by macroscopic techniques due to the nonaffine deformation of the densely cross-linked polymer network domains (Di Lorenzo et al., 2015). Indeed, the gelatin gels crosslinked with mTG are heterogenous due to the nature of enzymatic crosslinking, as well as the introduction of both intra-molecular or inter-molecular covalent bonds and steric factors (Yung et al., 2007). That was observed here as the larger coefficient of variation in the AFM mechanical maps over the gelatin gels in comparison with the PAAG and Ecoflex gels (Fig. 2). In another study (Mao et al., 2022), for similarly produced gelatin gels, the presence of a heterogeneous fiber network was shown by scanning electron microscopy imaging. Therefore, while AFM probes densely cross-linked network domains, since the involved length scale might be similar to that of these domains (a ∼100 nm contact radius between the tip and the sample), other used methods involve a much larger length scale and mostly measure the softer loosely cross-linked part of the gel (Di Lorenzo et al., 2015).

**Figure 6.**
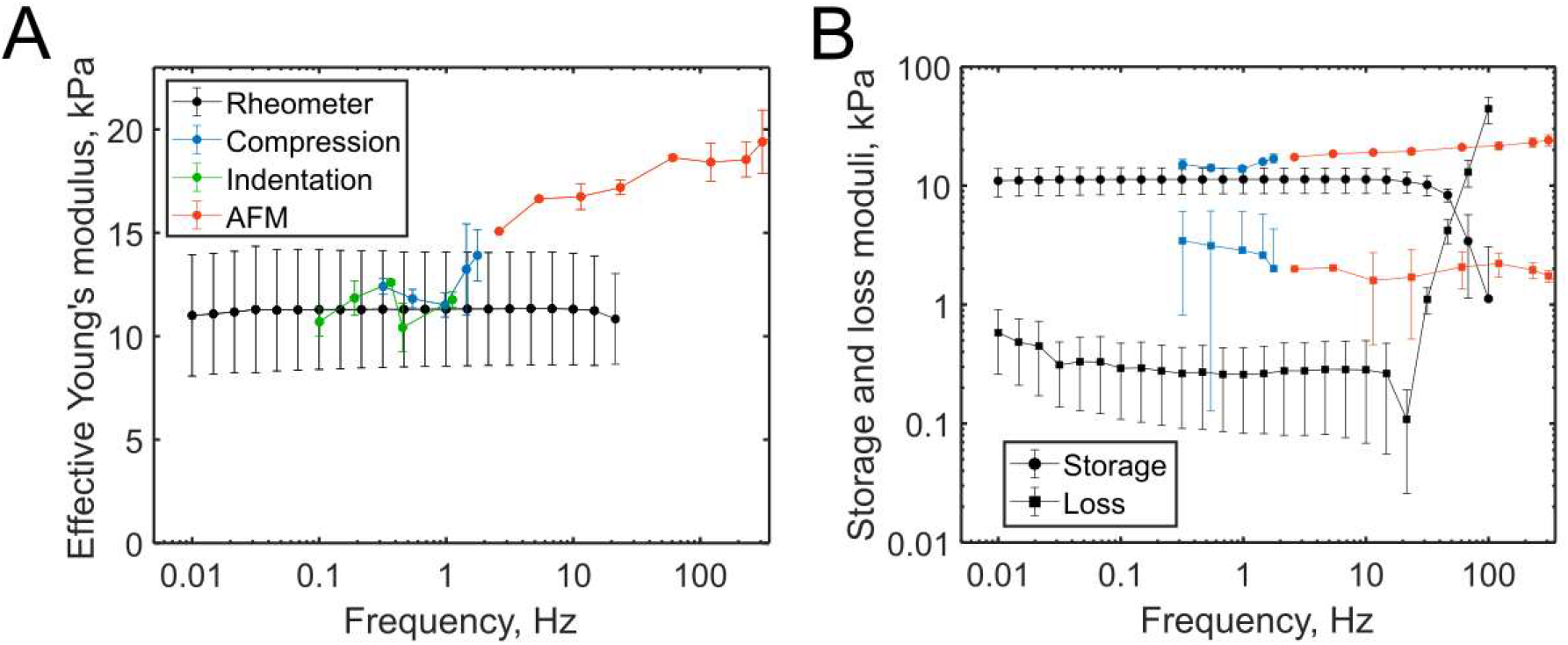
Comparison of five stiffness characterization methods for gelatin gels crosslinked with mTG. (A) Effective Young’s modulus versus effective frequency; (B) Storage and loss Young’s moduli versus effective frequency, microindentation data are not presented due to a too weakly expressed hysteresis in the indentation cycle. The data are presented as mean±SD for measurements on three samples.

### 3.4 Comparison of different techniques for mechanical characterization

The goal of this work was to evaluate whether different mechanical characterization methods for soft gels provide comparable data, especially those related to a frequency-dependent viscoelastic response. We characterized the mechanical parameters of gels from three different materials (PAAG, silicone Ecoflex, gelatin) using rheology, compression, microindentation, and nanoindentation. Overall, it might be concluded that the correspondence between the parameters provided by different techniques greatly depends on the studied material. For a homogenous linearly elastic material, such as PAAG, a good correspondence between all the methods was obtained. Even at high frequencies, when the viscoelasticity contribution started to be substantial, a good correlation between the methods with high available frequencies (rheology and AFM) was achieved.

For a material with a highly pronounced viscoelastic behavior, such as the Ecoflex silicone gel, it becomes most important to account for the characteristic frequencies applied during the measurements. Some techniques, such as compression and microindentation, are better adjusted for the low-frequency range, while such a technique as AFM is generally operated at higher frequencies. These characteristic frequencies can be a source of the divergence between the effective stiffness measured by the corresponding techniques, thus plotting the apparent modulus (or storage and loss moduli) versus the applied frequency is a more favorable method for the data comparison. It is especially important since many biological tissues are highly viscoelastic (Fung, 1993; Sasaki, 2012; Verdier et al., 2009). Heterogeneity is another property of biological materials which is expected to affect the measurements to different extents depending on the characteristic spatial scales.

Each of the applied methods has certain benefits, but also limitations and corrections which should be carefully accounted for (Griffin et al., 2016; Narasimhan et al., 2020; Navindaran et al., 2023; Arevalo et al., 2022). Rheology is beneficial because of the large available frequency range, but it requires sample dimensions being exactly adjusted to the measurement geometry. Slippage of the sample between the plates might occur at high frequencies (Herrada-Manchón et al., 2023; Megone et al., 2018).

Compression also requires samples with regular shapes and smooth surfaces, and the frequencies are generally more limited due to the system inertia. Microindentation is a promising technique due to less strict requirements for the sample, which may have an irregular shape but should be smooth at the spatial scale of the used indenter (Rubiano et al., 2019). The modern microindentation devices allow spatial mapping, selection of probes of different shapes and dimensions, and different experimental routines for viscoelasticity measurements (e.g., creep, force relaxation), and are often a part of universal mechanical measurement systems (Abba and Kalidindi, 2020). However, it is important to apply the required corrections for the processing, such as the correction for the finite sample thickness. Finally, AFM is a standard technique for the measurement of mechanical properties at the nano- and microscale, which means certain specificity related to this spatial scale, surface effects, and local heterogeneities (Cho et al., 2023; Efremov et al., 2020b). On the other hand, AFM provides the data at the scale where cells interact with the surrounding media, which is important in the development and testing of biomaterials.

## 4. Conclusions

In this study, we have examined the mechanical properties of three distinct types of soft gels commonly used in biomedical applications, employing four different mechanical testing techniques: shear rheology, compression, microindentation, and nanoindentation by atomic force microscopy. Notably, when analyzing an elastic homogeneous synthetic hydrogel (PAAG), all the methods provided closely aligned mechanical results. However, for a highly viscoelastic gel (such as Ecoflex) and a heterogeneous biopolymer hydrogel (gelatin crosslinked with mTG), the discrepancies between the methods were more pronounced. To ensure the accurate interpretation of the data, the majority of the methods required introduction of correction factors to account for the boundary conditions. Our findings underscore the importance of considering both the temporal and spatial scales inherent in each measurement technique. Furthermore, our study highlights the necessity of employing a combination of different techniques, particularly for heterogeneous viscoelastic materials such as biological samples. By elucidating these complexities, we contribute to the refinement of mechanical testing protocols and the advancement of soft gel design for biomedical applications, including bioprinting and development of tissue engineered constructs and bioequivalents.

## Acknowledgements

This work was supported by the Russian Science Foundation (grant No. 23-74-10113, https://rscf.ru/en/project/23-74-10113/, AFM, microindentation experiments and all data processing) and by the Ministry of Science and Higher Education of the Russian Federation within the framework of state support for the creation and development of World-Class Research Centers “Digital biodesign and personalized healthcare” №075-15-2022-304 (compression and rheological experiments).

